# *tombRaider* – improved species and haplotype recovery from metabarcoding data through artefact and pseudogene exclusion

**DOI:** 10.1101/2024.08.23.609468

**Authors:** Gert-Jan Jeunen, Kristen Fernandes, Eddy Dowle, Gracie C. Kroos, Quentin Mauvisseau, Michal Torma, Allison K. Miller, Miles Lamare, Neil Gemmell

**Affiliations:** Department of Marine Science, University of Otago, Dunedin 9016, New Zealand; Department of Anatomy, University of Otago, Dunedin 9016, New Zealand; Natural History Museum, University of Oslo, Oslo 0562, Norway; Department of Microbiology, University of Otago, Dunedin 9016, New Zealand

**Keywords:** environmental DNA, software, algorithm, PCR, amplification

## Abstract

Environmental DNA metabarcoding has revolutionized ecological surveys of natural systems. By amplifying and sequencing small gene fragments from environmental samples containing complex DNA mixtures, scientists are now capable of exploring biodiversity patterns across the tree of life in a time-efficient and cost-effective manner. However, the accuracy of species and haplotype identification can be compromised by sequence artefacts and pseudogenes. Despite various strategies developed over the years, effective removal of artefacts remains challenging and inconsistent data reporting standards hinder reproducibility in eDNA metabarcoding experiments. To address these issues, we introduce *tombRaider*, an open-source command line software program (https://github.com/gjeunen/tombRaider) and R package (https://github.com/gjeunen/tombRaider_R) to remove artefacts and pseudogenes from metabarcoding data post clustering and denoising. *tombRaider* features a modular algorithm capable of evaluating multiple criteria, including sequence similarity, co-occurrence patterns, taxonomic assignment, and the presence of stop codons. We validated *tombRaider* using various published data sets, including mock invertebrate communities, air eDNA from a zoo, and salmon haplotypes from aquatic eDNA. Our results demonstrate that *tombRaider* effectively removed a higher proportion of artefacts while retaining authentic sequences, thus enhancing the accuracy and reliability of eDNA-derived diversity metrics. This user-friendly software program not only improves data quality in eDNA metabarcoding studies, but also contributes to standardised reporting practices, an aspect currently lacking in this emerging research field.

## 1 INTRODUCTION

Biodiversity assessments are central to ecology and conservation, as they provide critical insights into ecosystem health, resilience, and function (Thorogood et al. 2023). Visual observation of species is often impractical for comprehensive quantification of multi-taxon biodiversity assessments, due to the complexity of biological systems and variability of life forms (Pereira et al. 2021). This hampers our ability to fully understand and protect ecosystems, highlighting the need for innovative methods and technologies (Ficetola et al. 2008; Thomsen and Willerslev 2015). Environmental DNA (eDNA) metabarcoding, i.e., the simultaneous identification of taxa from DNA found in environmental samples (Taberlet et al. 2012), has radically altered the cost-effectiveness and time-efficiency of large-scale biodiversity monitoring across the tree of life (Garlapati et al. 2019; Jeunen et al. 2024). Hence, ecological studies saw a substantial increase in the implementation of eDNA metabarcoding during the last decade (Beng and Corlett 2020; Takahashi et al. 2023).

Environmental DNA metabarcoding builds upon the premise that organismal genetic traces can be extracted from environmental samples, such as water (Jeunen et al. 2019), sediment (Xie et al. 2018), soil (Olmedo-Rojas et al. 2023), and air (Lynggaard et al. 2022). Due to the low concentration and complex genetic mixture contained within environmental samples, PCR (Polymerase Chain Reaction) amplification is carried out prior to high throughput sequencing to enrich samples for target barcoding genes holding interspecific variability for species detection (Stat et al. 2017) or intraspecific variability for population genetic analyses (Weitemier et al. 2021). Afterwards, raw sequence data is cleaned bioinformatically to generate a list of OTUs (Operational Taxonomic Units) or ASVs (Amplicon Sequence Variants) through clustering (Edgar 2010) and denoising (Edgar 2016b), respectively. These biologically-relevant sequences are then used as a proxy for species or haplotype detections to answer ecological questions (Antich et al. 2021). The ability of OTUs and ASVs to accurately reflect biological diversity is questioned, however, as metabarcoding frequently overestimates α-diversity (Frøslev et al. 2017; Andújar et al. 2021; Chiarello et al. 2022).

Many studies have observed an inflated number of OTUs and ASVs when compared to expected species richness (Peng et al. 2015; Zhou et al. 2019; Nagai et al. 2022). This issue became particularly apparent when transitioning from studies of microbial diversity to eukaryote monitoring, due to the generally better-understood species diversity of macroscopic organisms, e.g., fish, compared to the vast and often uncharacterised microbial communities within ecosystems. OTU/ASV inflation has been further explored through bacterial and eukaryotic mock communities (Brown et al. 2015; Flynn et al. 2015; Nagai et al. 2022), with increases observed as high as orders of magnitudes (Flynn et al. 2015). This inflation of α-diversity most likely stems from PCR and sequencing errors, henceforward referred to as “artefacts” (Frøslev et al. 2017), as well as the presence of nuclear mitochondrial pseudogenes (NUMTs) within environmental DNA extracts (Andújar et al. 2021).

Polymerases incorporate various mutations during deoxyribonucleotide chain synthesis, including recombination, transitions, transversions, insertions, and deletions (Zhou et al. 2019). The rate at which PCR errors are assimilated depends on the thermostability, fidelity, processivity, and specificity of the polymerase used (Nagai et al. 2022). The problem of PCR errors is further exacerbated in low-quantity and quality eDNA samples by the high number of PCR cycles needed for adequate amplification (Ishii and Fukui 2001; Kanagawa 2003), as well as primer sequence matching (Kumar et al. 2022), a known issue for metabarcoding primers amplifying broad taxonomic groups (Zhang et al. 2020). While chimeric sequences are successfully removed during bioinformatic analysis (Edgar et al. 2011), clustering (Edgar 2010) and denoising (Callahan et al. 2016; Edgar 2016b) algorithms are incorporated to account for minor differences between reads stemming from biological variation and PCR errors. However, the complexity of the eDNA signal and the high variability in starting abundances of species’ DNA hinders the algorithm’s ability to discriminate between PCR errors and genuine biological variants. This has led to the persistence of PCR errors in downstream statistical analyses, resulting in inflated α-diversity observations in metabarcoding biodiversity surveys (Frøslev et al. 2017). As PCR artefacts are unique and, therefore, sample specific, their presence can also influence observed β-diversity patterns.

Environmental DNA metabarcoding surveys are susceptible to further inflated α-diversity, due to the co-amplification of NUMTs (Andújar et al. 2021). Over time, NUMTs accumulate within host nuclear genomes and can undergo duplication events to form NUMT families (Pamilo et al. 2007; Baldo et al. 2011). The inclusion of functional mitochondrial sequences within NUMTs, particularly those integrated into invariant regions of the nuclear genome (Bensasson et al. 2001), increases the likelihood of co-amplification during PCR. This issue is exacerbated by wobble (degenerate) bases, which are frequently incorporated in metabarcoding primers to capture the known mitochondrial variation in primer-binding regions across the taxonomic group of interest (Leray et al. 2013).

In an attempt to obtain more accurate α- and β-diversity metrics from eDNA metabarcoding data, multiple strategies have been proposed to remove persistent artefacts and pseudogenes post bioinformatic processing. Arbitrary and estimated abundance thresholds are frequently used to discard spurious OTUs/ASVs representing PCR errors created during the latter stages of amplification (Brown et al. 2015; Bálint et al. 2016). However, PCR errors created early in the PCR process will remain, as these erroneous templates would occur in large copy numbers (Pienaar et al. 2006). Furthermore, abundance thresholds might discard genuine biological entities, with taxon read count being heavily influenced by primer-template binding affinities (Krehenwinkel et al. 2017; Liu et al. 2023). Locating the presence of stop codons to identify pseudogenes, on the other hand, is restricted to protein-coding genes and fails where mutations do not amount to stop codons (Andújar et al. 2021). Merging OTUs/ASVs based on their taxonomic ID, a frequently-used method in eukaryotic metabarcoding studies, has the potential to combine sequences that should be kept separate when closely related species are assigned identical taxonomic ID’s due to misclassification resulting from incomplete reference databases (Keck et al. 2023; Marinchel et al. 2023) and is not appropriate when haplotype data is of value. Finally, the identification of artefacts based on co-occurrence patterns and a fixed threshold for sequence similarity (Frøslev et al. 2017) has been shown to merge co-occurring congeners into a single OTU/ASV. Thus, although metabarcoding is known to artificially inflate α-diversity through the incorporation of PCR errors and co-amplification of pseudogenes (Frøslev et al. 2017; Andújar et al. 2021), our current strategies for dealing with this issue are inadequate and potentially biased. Data curation to remove artefacts is further complicated by the inconsistent use of terminology and lack of standardised analysis reporting in eDNA research. An easy-to-use software program able to more accurately distinguish artefacts from genuine biological entities would aid metabarcoding biomonitoring to obtain reliable α- and β-diversity metrics without losing “real” signals, as well as enable standardised data analysis reporting.

Here, we introduce *tombRaider*, an open-source software package for improved species and haplotype recovery from metabarcoding data through accurate artefact and pseudogene exclusion. *tombRaider* has been developed to meet an unmet need regarding eDNA data analysis and reporting that we identified through a meta-analysis of 539 metabarcoding papers published in open-access journals during 2023. To determine *tombRaider’s* efficiency, we compared the expected diversity from a previously-published mock community (Braukmann et al. 2019) with the diversity observed using various data curation strategies. Additionally, we show *tombRaider’s* ability to uncover so-called relics, i.e., genuine biodiversity missed in previously-published data sets, based on zoo animals from air eDNA (Lynggaard et al. 2022). Finally, we provide evidence for *tombRaider* to recover haplotype diversity from metabarcoding data using a previously-published salmon eDNA haplotype study (Weitemier et al. 2021). We show that *tombRaider* provides non-significant differences in α- and β-diversity with expected results from mock communities, as well as being able to identify relics from past projects, and successfully identify haplotype diversity of multiple species simultaneously. Additionally, *tombRaider* is feature-rich, highly versatile in its implementation, and requires relatively limited computational resources.

## 2 METHODS

### 2.1 META-ANALYSIS

A systematic literature search was conducted to identify publications using metabarcoding methodologies. Peer-reviewed manuscripts published between the 1^st^ of January 2023 and the 10^th^ of November 2023 containing “metabarcoding” in the title, abstract, or keywords were searched using the Scopus citation database on the 10^th^ of November 2023. We restricted our search to open-access journal articles to limit the number of publications and excluded conference proceedings, book chapters, and commentaries/notes. We also limited our search to articles written in English. We randomly sub-selected 600 out of 771 publications to further reduce the number of included articles. We screened all publications individually to determine if methodologies employed metabarcoding techniques using the Illumina sequencing platform. Publications not meeting all requirements were excluded, with 539 publications remaining for the meta-analysis (Supplement 1). We recorded the following information for each of these publications: (i) the choice of clustering or denoising and program used, (ii) the post-clustering data curation method and program used, and (iii) sequence data and code availability. To show overall trends in data analysis across publications, an alluvial diagram was created using RAWGraphs *v* 2.0 (Mauri et al. 2017).

### 2.2 THE SOFTWARE PROGRAM

*tombRaider* is a versatile, open-source software program to remove artefacts and pseudogenes from metabarcoding datasets. *tombRaider* is available on GitHub (https://github.com/gjeunen/tombRaider) and coded in base Python 3 through dynamic programming to limit the number of dependencies and simplify installation. An R wrapper is also available (https://github.com/gjeunen/tombRaider_R) to support bioinformatic pipelines conducted in the R environment, e.g., DADA2 (Callahan et al. 2016). *tombRaider* features a modular algorithm capable of evaluating a range of criteria individually or in combination, including (i) sequence similarity, (ii) co-occurrence patterns, (iii) taxonomic assignment, and (iv) the presence of stop codons. This flexibility enables users to define which criteria are incorporated in the identification of artefacts, ranging from the investigation of a single criterion (e.g., taxonomic assignment: merging sequences based on taxonomic ID) to a combination of multiple criteria (e.g., sequence similarity – co-occurrence patterns: LULU (Frøslev et al. 2017)). *tombRaider*’s workflow (Figure 1) is contained within a single line of code for user-friendliness, whereby *tombRaider* imports three mandatory input files that are generated during standard bioinformatic processing, as well as an optional alignment file to increase execution speed. After importing the data, *tombRaider* runs the algorithm, and outputs updated documents from which artefacts are removed, as well as a simple log file for standardised data reporting.

**Figure 1:**
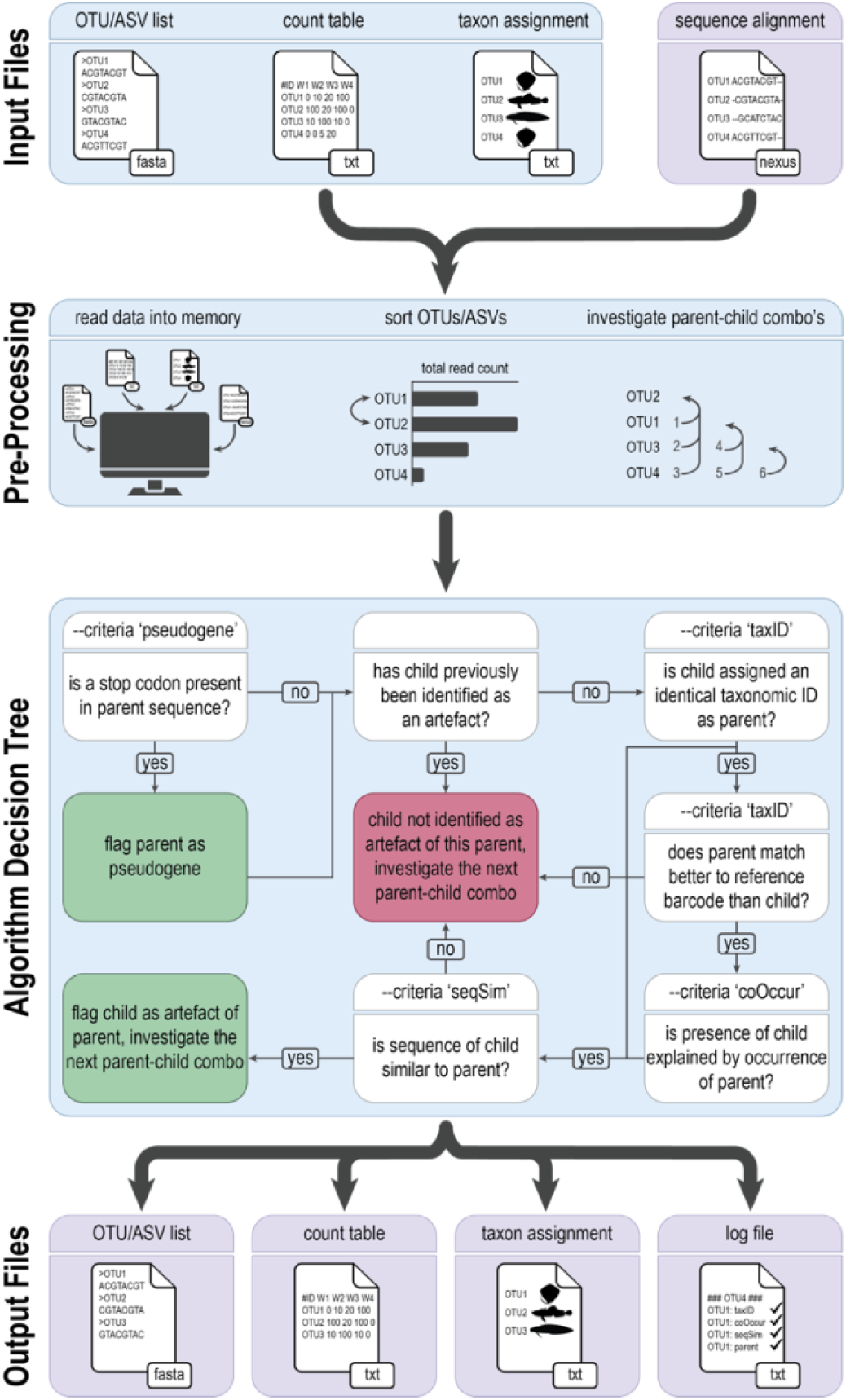
tombRaider’s workflow. Users are required to provide three mandatory files, while tombRaider can also import an optional alignment file to speed up code execution. Once files are read into memory and parameters checked, tombRaider sorts OTUs/ASVs based on the ‘--sort’ parameter. Next, all parent (highest order) – child (lowest order) combinations will be checked sequentially by the algorithm decision tree to flag OTUs/ASVs either as pseudogene (‘--criteria pseudogen’) or PCR artefact. The algorithm decision tree will be dependent on the user-provided list of criteria. OTUs/ASVs flagged as pseudogene or artefact will be removed from the data prior to writing the results to output files.

#### 2.2.1 INPUT FILES

*tombRaider* requires three input files that are accessible to users upon completion of standard bioinformatic pipelines. The first input file is a tab-delimited OTU/ASV table (‘*--frequency-input*’). *tombRaider* expects samples as columns and OTUs/ASVs as rows, with the option for users to transpose the table if needed by providing the ‘*--transpose*’flag. To exclude metadata columns and rows from the table, users can list column headers and row names using the ‘*--omit-columns*’ and ‘*--omit-rows*’ parameters, respectively. Finally, samples such as negative and positive controls can be excluded from the analysis through ‘*--exclude*’ by specifying a list of sample names (column headers). The second input file is an OTU/ASV sequence file in a two- or multi-line fasta format (‘*--sequence-input*’), whereby sequence headers match the row names of the OTU/ASV table. The third, and final, required input file is a taxonomy assignment file (‘*--taxonomy-input*’). *tombRaider* currently supports four taxonomic classifiers, including BLAST (Altschul et al. 1990), BOLD (Ratnasingham and Hebert 2007), SINTAX (Edgar 2016a), and IDTAXA (Murali et al. 2018). *tombRaider* automatically identifies the classifier used based on the file format provided to ‘*--taxonomy-input*’. Additional parameters are needed dependent on the taxonomic classifier. For example, the ‘*--blast-format*’ parameter specifies the information within each column of the input file. Alternatively, ‘*--bold-format*’ specifies if a summary table was downloaded from the website (‘*--bold-format* summary’) or if a BOLDigger output file (‘*--bold-format* completÈ) was provided (Buchner and Leese 2020). Finally, the ‘*--sintax-threshold*’ parameter can be flagged when users prefer to use the column within the SINTAX input file containing the user-defined identity threshold. A multiple sequence alignment file in nexus format can optionally be imported into *tombRaider* using the ‘*--alignment-input*’ parameter. Importing a multiple sequence alignment file will circumvent *tombRaider* to calculate pairwise sequence alignments to investigate sequence similarity between OTU/ASV combinations and, thereby, significantly increasing the speed of code execution. Examples of each file type are available on the GitHub repository to aid users with formatting.

#### 2.2.2 THE ALGORITHM

Upon importing the input files, users can reorder the ‘*--frequency-input*’ table using the ‘*--sort*’ parameter. OTUs/ASVs can be reordered based on total read count (‘*--sort* “total read count”’), average read count (‘*--sort* “average read count”’), or detection frequency (‘*--sort* detections’). The algorithm will then sequentially investigate all lower-ranked (henceforward referred to as “child”) to higher-ranked OTUs/ASVs (henceforward referred to as “parent”). The criteria investigated by the algorithm depend on the user-specified list within the ‘*--criteria*’ parameter (Figure 1). When all user-specified criteria hold true, the child is flagged an artefact of the parent. Once an OTU/ASV is deemed an artefact, it will not be further assessed as a child for other potential parents, though can be found a parent itself for lower-abundant OTUs/ASVs. NUMTs are flagged separately within this sequential investigation of parent-child combos. Once all combinations are checked, artefact reads are summed to their identified parent prior to removal of the artefact (child – parent merging), unless the parent was previously deemed an artefact itself, in which case the artefact reads are summed to the highest-ranking OTU/ASV not deemed an artefact instead (child – grandparent merging). Alternatively, users can opt to discard artefacts outright without summing reads to the higher-ranked OTUs/ASVs by specifying the ‘*--discard-artefacts*’ flag. Finally, OTUs/ASVs flagged as pseudogenes are discarded from the data.

#### 2.2.3 USER-DEFINED PARAMETERS

Users can provide additional parameters and set thresholds to identify artefacts. To reduce the impact of spurious detections from tag jumping (Schnell et al. 2015), an abundance threshold can be set for what *tombRaider* considers a “true” detection (‘*--detection-threshold*’). Additionally, the sequence similarity threshold can be specified using the ‘*--similarity*’ parameter. Sequence similarity between parent and child can either be calculated by *tombRaider* through native implementations of the Needleman-Wunsch (Needleman and Wunsch 1970) (‘*--pairwise-alignment* “global”’) and Smith-Waterman (Smith and Waterman 1981) algorithms (‘*--pairwise-alignment* “local”’) or inferred from the genetic distance based on the optional multiple sequence alignment input file (‘*--alignment-input*’). Furthermore, users can specify if the co-occurrence pattern should be assessed on a presence-absence transformed data frame (‘*--occurrence-type* “presence-absence”’) or based on read abundance (‘*--occurrence-type* “abundance”’), as well as the ratio type and threshold of the co-occurrence pattern (‘*--occurrence-ratio*’), including (i) the minimum ratio across all samples (‘*--occurrence-ratio* “global;1.0”’), (ii) the minimum ratio across all samples with positive detections (‘*--occurrence-ratio* “local;1.0”’), or (iii) the maximum number of samples where a child is present without the parent (‘*--occurrence-ratio* “count;1”’). Users can also set the ‘*--taxon-quality*’ flag, which requires a child not only to be assigned to the same taxonomic ID as a potential parent, but also for the parent to obtain a better or equal match to the reference sequence compared to the child. Finally, when users are interested to explore intraspecific variation, the taxonomic ID can be switched to the reference sequence ID by setting the ‘*--use-accession-id*’ flag. This option is only available when BLAST (Altschul et al. 1990) is used as taxonomic classifier, as the remaining classifiers do not provide information on the reference sequence an ASV/OTU matched against.

#### 2.2.4 OUTPUT FILES

Upon completion of the analysis, *tombRaider* will write updated files without artefacts for each input file when ‘*--frequency-output*’, ‘*--sequence-output*’, and ‘*--taxonomy-output*’ are specified. These output files are formatted as the input files and can be imported into any statistical analysis pipeline, such as the phyloseq R package (McMurdie and Holmes 2013). Additionally, a log output file (‘*--log*’) can be generated for closer inspection on why sequences were deemed artefacts or true ASVs/OTUs.

### 2.3 ALGORITHM VALIDATION

We validated *tombRaider* using various published data sets, including invertebrate mock communities (Braukmann et al. 2019), air eDNA from the Copenhagen Zoo (Lynggaard et al. 2022), and salmonid haplotypes from US streams (Weitemier et al. 2021). To verify *tombRaider*’s compatibility with all main bioinformatic pipelines, published datasets were bioinformatically processed prior to data curation using a variety of frequently-utilised software programs in metabarcoding research, including DADA2 (Callahan et al. 2016), USEARCH/VSEARCH (Edgar 2010; Rognes et al. 2016), and QIIME (Caporaso et al. 2010). A full description of the bioinformatic and statistical analyses for each dataset can be found in Supplement 2 (invertebrate mock communities), Supplement 3 (air eDNA from the Copenhagen Zoo), and Supplement 4 (salmonid haplotypes from US streams), respectively. All bioinformatic and statistical analysis scripts, on the other hand, can be retrieved from the following GitHub repository: https://github.com/gjeunen/tombRaider_supplemental_files.

#### 2.3.1 MOCK COMMUNITY

The terrestrial arthropod mock communities were comprised of 367 unique taxa, each constructed with different body segments (abdomen, bulk leg, composite leg) and slightly altered species compositions (Braukmann et al. 2019). Illumina sequencing data was downloaded from NCBI’s Short Read Archive (SRP158933) and bioinformatically processed using a standard cutadapt *v* 4.4 (Martin 2011) and VSEARCH *v* 2.16.0 (Rognes et al. 2016) denoising pipeline. ASV sequences, a ∼410 bp fragment of the cytochrome c oxidase subunit I gene, were assigned a taxonomic ID through BOLDigger (Buchner and Leese 2020). Phylogenetic α-(Faith’s PD) and β-diversity (unweighted UniFrac distance) metrics were calculated for a variety of data curation treatments and compared against the known mock community composition to assess the importance of criteria in identifying artefacts, including (i) unfiltered (no data curation post denoising), (ii) taxon-dependent co-occurrence merging (criteria evaluated: taxonomic ID, sequence similarity, co-occurrence pattern), (iii) taxonomic ID merging (criteria evaluated: taxonomic ID), (iv) taxon-independent co-occurrence merging (criteria evaluated: sequence similarity, co-occurrence pattern), and (v) abundance filtering.

#### 2.3.2 REANALYSING PUBLISHED DATA TO RECOVER RELICS

To determine *tombRaider’s* capability in recovering relics, i.e., species and haplotypes missed during initial analysis, from published data sets, we reanalysed Illumina sequence data from the Copenhagen Zoo air eDNA experiment (Lynggaard et al. 2022). Sequences for two amplicons (Taylor 1996; Riaz et al. 2011) were downloaded from the University of Copenhagen Electronic Research Archive and processed via the DADA2 *v* 1.26 (Callahan et al. 2016) denoising platform. Custom reference databases were generated for each primer set using CRABS *v* 0.1.8 (Jeunen et al. 2022) for taxonomic assignment of ASVs using local blastn *v* 2.10.1+ searches (Altschul et al. 1990). Species detections after data curation with *tombRaider* were compared against the animal inventory list of the Copenhagen Zoo, as well as the species list provided by Lynggaard et al. (2022) for which sequence data was clustered into OTUs and curated by LULU.

#### 2.3.3 SALMONID HAPLOTYPES

To investigate the potential of *tombRaider* to recover intra-specific variation, i.e., haplotypes, from metabarcoding data, we reanalysed the NADH dehydrogenase 2 (ND2) dataset from (Weitemier et al. 2021) for three salmonids simultaneously, including *Oncorhynchus clarkii clarkii* (Richardson, 1836), *O. kisutch* (Walbaum, 1792), and *O. tshawytscha* (Walbaum, 1972). The ASV table and sequences were obtained from the online Supplementary Files associated with Weitemier et al. (2021). A highly curated salmonid reference database was generated for the ND2 gene using CRABS *v* 0.1.8 (Jeunen et al. 2022) prior to conducting a local blastn *v* 2.10.1+ search (Altschul et al. 1990) to assign a taxonomic ID to each ASV. Haplotype networks were generated for each salmonid independently using the R packages pegas *v* 1.3 (Paradis 2010) and adegenet *v* 2.1.10 (Jombart 2008). *tombRaider* was run to identify previously observed haplotypes (i.e., ASVs with a perfect match to the reference database), potentially novel haplotypes (i.e., ASVs deemed true, but without a perfect match to the reference database), and artefact sequences. The *tombRaider* output was finally incorporated into the haplotype networks to visualize *tombRaider*’s efficiency to recover intra-specific variation from metabarcoding data.

## 3 RESULTS

### 3.1 META-ANALYSIS

We identified 539 publications that matched the search criteria for the meta-analysis, of which 533 had adequate methodology and/or supplementary files to be assessed. Across all publications, there was a roughly 60/40 split between denoising (ASVs; 321 publications) and clustering (OTUs; 212 publications). Denoising methodologies included publications that had stated they used zero-distance OTUs (ZOTUs) or exact sequence variants (ESVs). For ease of presentation, we have categorised these into ‘ASVs’. A wide range of software pipelines were used to process data (Figure 2). Predominantly for denoising methods, DADA2 (Callahan et al. 2016) was used, but for clustering, these methodologies varied. The majority of publications did not conduct data curation post clustering. Though for the ones that did, merging based on taxonomic ID (26%) was most-frequently reported. However, for 89 publications (16%), it was unclear whether sequence clusters were curated post-generation (Figure 2). To confirm that bioinformatic methodologies were categorised correctly and to clarify where methods were unclear, we examined the ASV/OTU tables generated by each study and bioinformatic codes supplied by the authors. However, an alarming trend emerged, whereby most publications did not make ASV/OTU tables available (72.5%) nor the bioinformatic and statistical scripts (81.1%) (Figure 2). We considered instances where this information was ‘Available on request’ from authors as not available for the purpose of this study. Additionally, we did not check the validity of the provided links to data and script repositories when listed in the publication.

**Figure 2:**
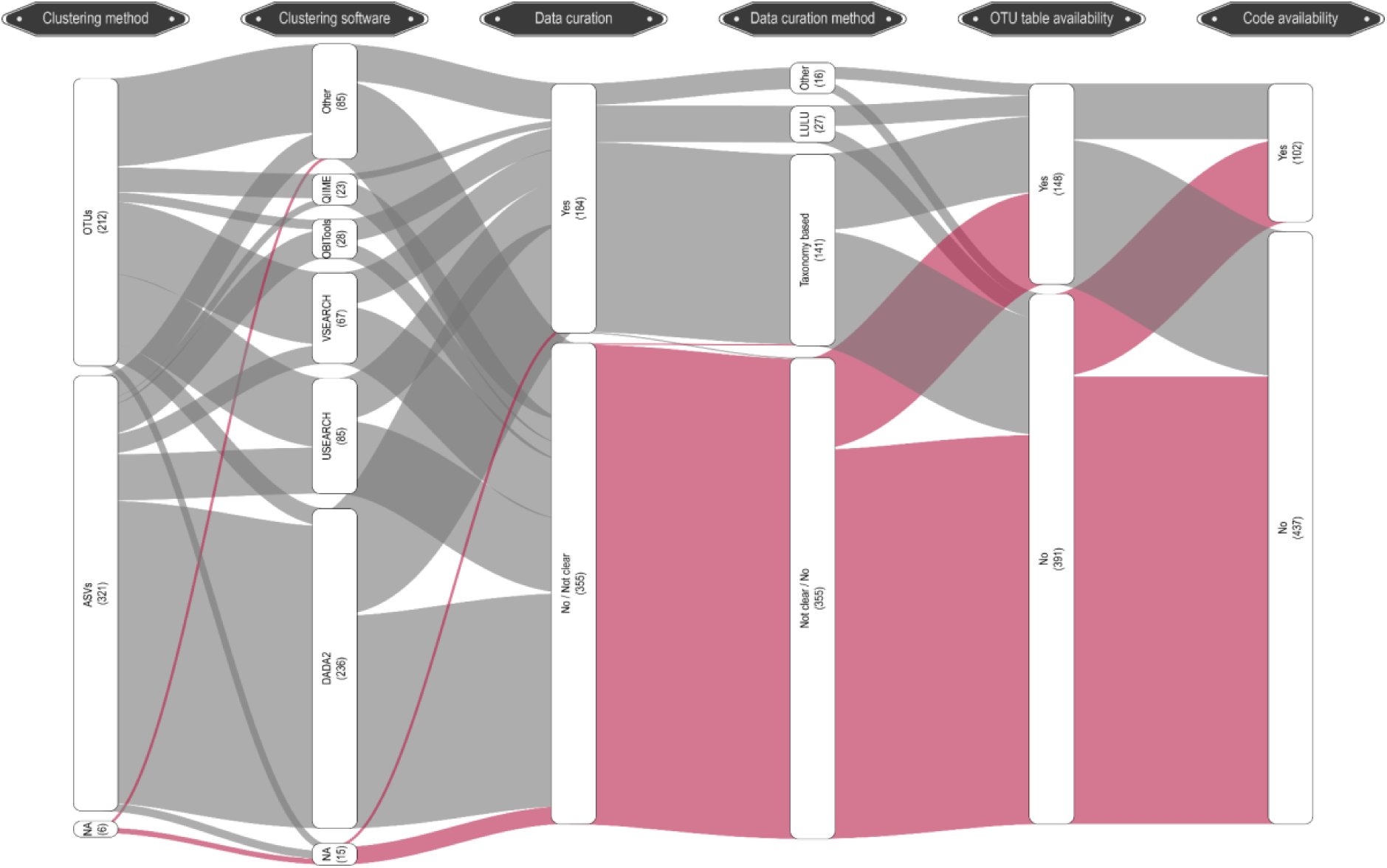
An alluvial diagram showing the number of publications analysed from our literature search. This diagram shows the clustering/denoising methodology used, the software program used for this process, whether or not clusters were merged, the software program/technique used for merging, OTU table availability, and code availability. Numbers in between brackets represent the number of publications. Publications with insufficient/missing information are indicated in pink.

### 3.2 INVERTEBRATE MOCK COMMUNITY

Bioinformatic processing of the invertebrate mock communities containing 367 unique terrestrial arthropod taxa resulted in a total of 798 ASVs (Figure 3a). Taxonomic assignment of the ASVs prior to data curation identified 337 mock community taxa and 30 taxa missing from the metabarcoding dataset. Additionally, 36 ASVs were identified as “potential contaminant”, whereby a perfect match was achieved for a species not in the mock community, leaving a total of 425 sequences identified as “artefact”. We obtained a dataset most closely resembling the mock community for the taxon-dependent co-occurrence merging treatment (mock community: 336; missing: 31; contaminant: 35; artefacts: 51), followed by taxonomic ID merging (mock community: 329; missing: 38; contaminant: 33; artefacts: 31), abundance filtering (mock community: 321; missing: 46; contaminant: 17; artefacts: 128), and taxon-independent co-occurrence merging (mock community: 127; missing: 240; contaminant: 10; artefacts: 12) (Figure 3b). While all data curation methods increased the distribution of the sequence similarity scores for retained sequences (Figure 3c), ANOVA (*F*_5,48_ = 2684, *p* < 0.0001***) and *post hoc* Tukey HSD revealed taxon-dependent co-occurrence merging as the only data curation method not significantly differing from the expected phylogenetic α-diversity measure (Faith’s PD) of the mock community (Figure 3d). Furthermore, visualization of phylogenetic β-diversity (unweighted UniFrac) showed theoretical mock community samples overlapping with the taxon-dependent co-occurrence merging and taxonomic ID merging treatments. Abundance filtering and unfiltered data, on the other hand, separated on the secondary axis explaining 19.9% of the variation within the dataset, while the taxon-independent co-occurrence merging separated on the primary axis explaining 74.8% (Figure 3e).

**Figure 3:**
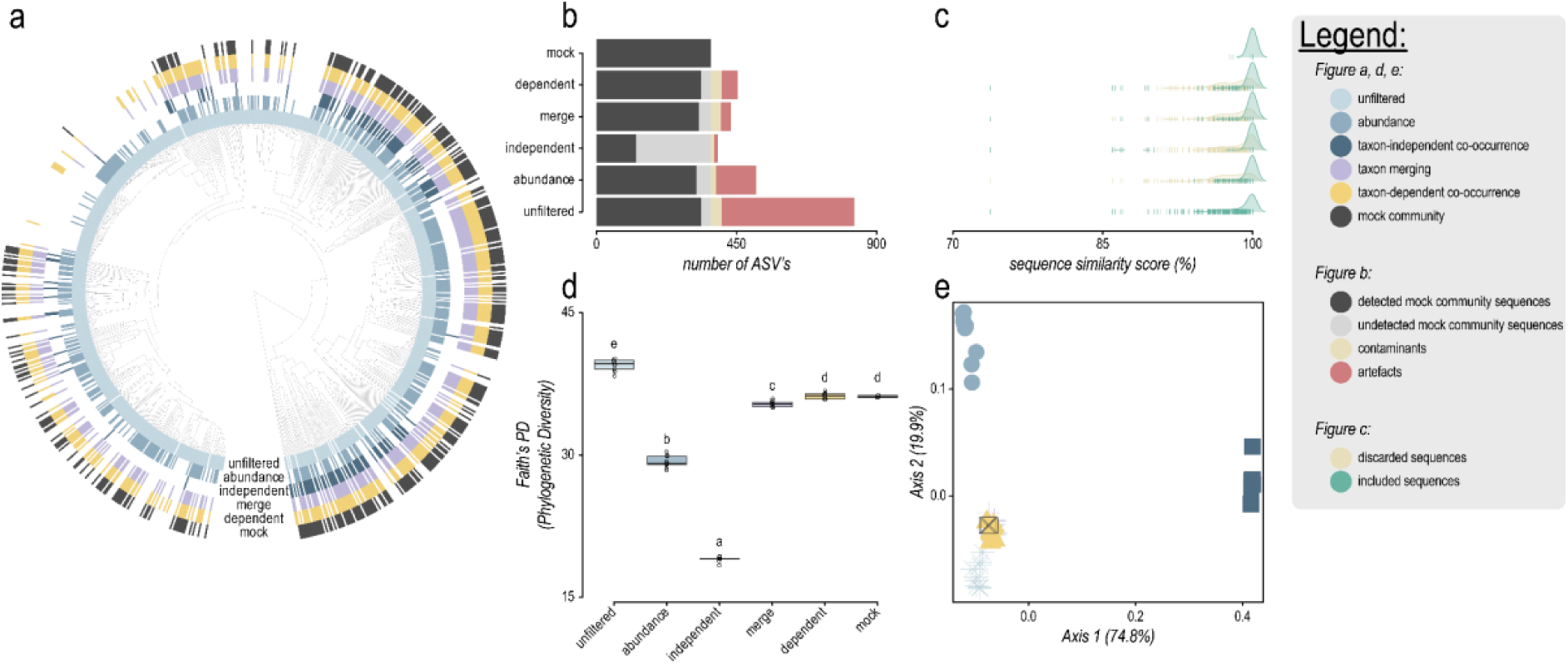
Invertebrate mock community analysis. (a) Bayesian phylogenetic tree with detections indicated by a coloured bar for the different data curation methods, including mock community species (black), taxon-dependent co-occurrence merging (“dependent”; yellow), taxonomic ID merging (“merge”; purple), taxon-independent co-occurrence merging (“independent”; dark blue), abundance filtering (“abundance”; blue), and unfiltered (light blue). (b) Number of ASVs detected by each method, with mock community species in black, missed mock community species in grey, potential contaminants (ASVs with a perfect match to a species not indicated as a mock community species) in gold, and sequence artefacts in red. (c) Distribution of sequence similarity scores for ASVs included after data curation (green) and discarded ASVs (gold). (d) Boxplots representing phylogenetic α-diversity comparison (Faith’s PD) between different methods. Letter represents statistically significant differences between treatments according to one-way ANOVA and post-hoc Tukey HSD. (e) Ordination plot depicting phylogenetic β-diversity (unweighted Unifrac distance) differences between treatments.

### 3.3 AIR EDNA

The DADA2 bioinformatic pipeline recovered 793 and 392 ASVs for the 12SV05 (Riaz et al. 2011) and 16Smam (Taylor 1996) primer sets, respectively. Data curation with *tombRaider* identified 258 (32.53%) and 96 (24.49%) ASVs as artefacts when considering the taxonomic ID, sequence similarity, and co-occurrence pattern criteria. The Copenhagen Zoo counted a total of 184 Chordates during sample collection, including 17 Amphibia, 67 Aves, 4 Actinopterygii, 63 Mammalia, and 33 Lepidosauria (Supplement 5). Using a taxon-independent co-occurrence merging approach (i.e., LULU), Lynggaard et al. (2022) reported the detection of 39 (21.2%) zoo animals. Reanalysis through *tombRaider*, on the other hand, detected an additional 50 (27.17%) species, bringing the total of detected animals kept in the Copenhagen Zoo to 89 (47.83%), including 3 (17.65%) amphibians, 36 (53.73%) birds, 0 (0%) fish, 48 (76.2%) mammals, and 2 (6.06%) reptiles (Figure 4).

**Figure 4:**
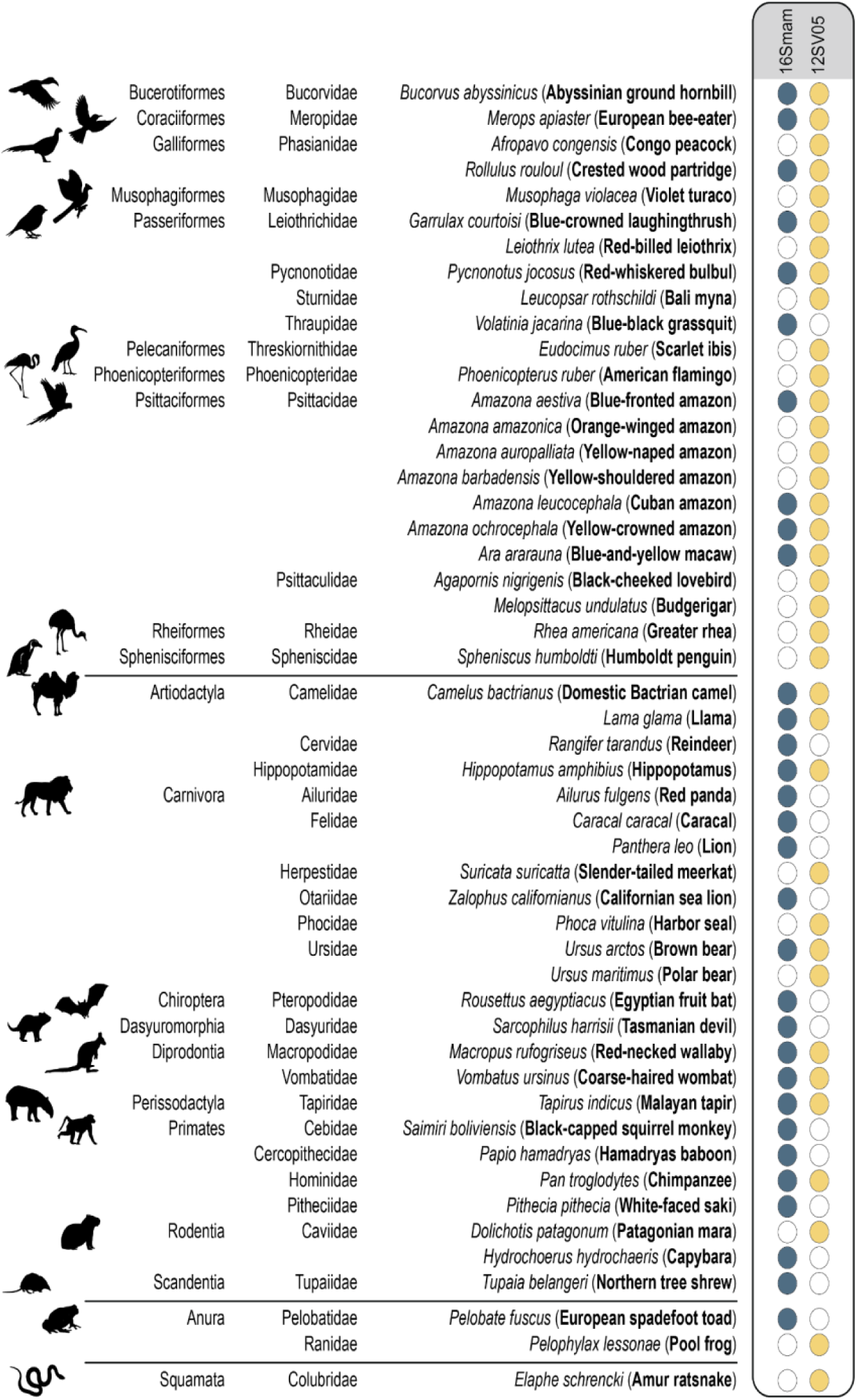
List of Copenhagen Zoo animals additionally detected from air eDNA through re-analysis with tombRaider. Taxa detected by the initial Lynggaard et al. (2022) analysis are omitted from the figure. Taxonomic order and family are listed for each species, with common names provided in between brackets in bold. Coloured circles denote for which primer set a species was detected, including 16Smam in blue (Taylor 1996) and 12SV05 in yellow (Riaz et al. 2011). Silhouettes for each order are provided on the left-hand side. Species are split up between classes using a solid black line. From top to bottom: Aves, Mammalia, Amphibia, Lepidosauria.

### 3.4 SALMONID HAPLOTYPES

The initial data (Weitemier et al. 2021) reported a total of 40 ASV’s as potential salmonid haplotypes, with 13 ASV’s for *O. clarkii*, 16 ASV’s for *O. kisutch*, and 11 ASV’s for *O. tshawytscha*. Of these potential haplotypes, eight ASV’s (20%) matched perfectly to a reference sequence in the database, including five *O. clarkii* ASV’s (38.46%), two *O. kisutch* ASV’s (12.50%), and one *O. tshawytscha* ASV (9.09%). A phylogenetic analysis containing the 40 ASV’s and all 64 currently available haplotype reference sequences for the three salmonids (*O. clarkii*: 56; *O. kisutch*: 4; *O. tshawytscha*: 4) revealed monophyletic groups for each species, as well as visualised the importance of artefact removal prior to statistical analysis (Figure 5). *tombRaider* successfully retained all eight ASV’s matching perfectly to a reference sequence, as well as identified an additional four ASV’s as potentially novel haplotypes and the remaining 28 ASV’s as likely artefacts introduced during laboratory processing of eDNA samples (Figure 5).

**Figure 5:**
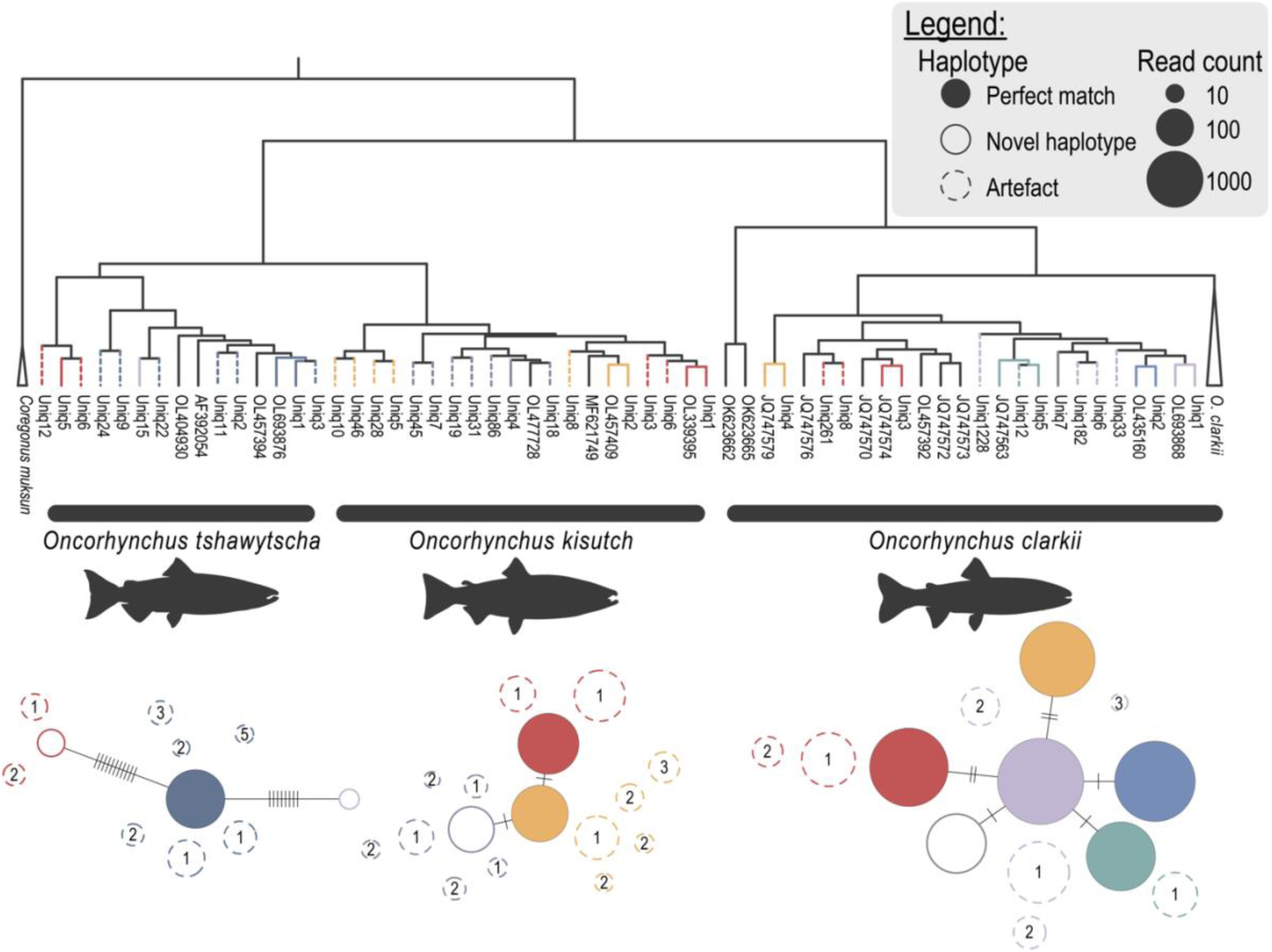
Salmonid haplotype analysis. Phylogenetic tree and haplotype networks of three salmonid species obtained from aquatic eDNA metabarcoding of the NADH dehydrogenase 2 gene. The phylogenetic tree consists of ASV’s, as well as currently available reference sequences. Branches are coloured based on tombRaider results, with identified haplotypes coloured similarly as their reference sequence match and artefacts represented as dotted lines in the same colour. Tip labels represent NCBI accession numbers for reference sequences or ASV identifiers for metabarcoding data. Within the haplotype networks, observed ASVs matching previously sequenced haplotypes are represented as filled circles. Potentially novel haplotypes, i.e., ASVs not perfectly matching a previously sequenced haplotype are represented as outlined circles. ASVs identified by tombRaider as artefacts are displayed as outlined circles with a dotted line. Haplotype colours match colours of the corresponding phylogenetic tree branches. Artefact colours match the haplotype it was merged with. Circle size represents the total number of reads associated with ASV across all samples. Number within artefacts represent the number of mismatches with the merged sequence.

## 4 DISCUSSION

The need for additional data curation to remove persistent artefacts and NUMTs after bioinformatic analysis and enable reliable α- and β-diversity metrics from metabarcoding data is a known, but understudied issue (Frøslev et al. 2017; Andújar et al. 2021). With an exponential increase in the implementation of metabarcoding for biodiversity monitoring across the globe, it is imperative simple approaches to remove artefacts and NUMTs from data sets are available for end-users. Here, we present *tombRaider*, an easy-to-use software program featuring a modular algorithm capable of accurately identifying and removing artefacts and NUMTs based on a user-defined list of criteria. The modularity of the algorithm enables flexibility for users to choose which criteria to include for identifying artefacts, while also allowing for easy incorporation of novel criteria in the future upon advancements in the metabarcoding research field. Furthermore, this modular approach provides users the opportunity to specify multiple criteria to be assessed for the identification of artefacts, thereby eliminating the specific biases associated with each criterion and obtain reliable α- and β-diversity metrics from metabarcoding data. *tombRaider* also facilitates standardized data reporting by generating a log file containing all necessary metadata for reproducibility of the results, an aspect currently lacking in this emerging research field. *tombRaider* is available as a command-line program (https://github.com/gjeunen/tombRaider), as well as an R package (https://github.com/gjeunen/tombRaider_R) to facilitate easy incorporation into existing bioinformatic pipelines.

We detected more than double the number of expected ASVs based on the species in a previously published mock community (Braukmann et al. 2019). This level of OTU/ASV inflation, despite stringent quality filtering during bioinformatic processing, aligns with findings from other eDNA metabarcoding studies (Flynn et al. 2015; Nagai et al. 2022). The significant differences in α- and β-diversity due to the increased number of ASVs highlight the critical need for data curation post bioinformatic analysis to obtain reliable metrics from eDNA metabarcoding data. Prior research has linked OTU/ASV inflation to PCR and sequencing artefacts (Flynn et al. 2015; Frøslev et al. 2017; Zhou et al. 2019) and the co-amplification of NUMTs when targeting protein-coding genes (Andújar et al. 2021). While considerable effort has been made to investigate the impact of polymerases on artefact proportion (Nagai et al. 2022), effective data curation remains essential until improvements in PCR fidelity are achieved (Gilje et al. 2008; Filges et al. 2019) or metabarcoding workflows incorporate non-PCR-based techniques for target enrichment and library preparation (Wilcox et al. 2018; Li et al. 2023). Accurate identification of artefacts remains challenging, as evidenced by significant discrepancies between current data curation methods and the known mock community composition, including abundance filtering (Drake et al. 2022), artefacts identification based on sequence similarity and co-occurrence patterns (Frøslev et al. 2017), and merging ASVs with identical taxonomic IDs.

The modular algorithm of *tombRaider* allows users to specify multiple criteria to distinguish true biological entities from artefacts. This capability led to the taxon-dependent co-occurrence merging data curation approach as being the only method in our mock community analysis (Braukmann et al. 2019) that did not show significant deviations from expected α- and β-diversity metrics. This enhanced accuracy is likely due to the algorithm’s decision tree incorporating a greater number of criteria, which helps mitigate the unique biases associated with each criterion when used in isolation. Additionally, by more effectively identifying artefacts, *tombRaider* facilitates the recovery of lost relics, as evidenced by a 178% increase in detected zoo animals (Lynggaard et al. 2022) when reanalysing previously published datasets. Moreover, the algorithm’s ability to accurately assess haplotype diversity across multiple species simultaneously (Weitemier et al. 2021) promises to advance the field of eDNA metabarcoding from species detection to evaluating population genetic structures and gene flow, by exploring intraspecific variability (Andres et al. 2023; Couton et al. 2023; Liu et al. 2024).

The artefact identification method based on co-occurrence patterns of similar sequences, known as LULU and pioneered by Frøslev et al. (2017), exhibited the most pronounced deviation from the expected phylogenetic α- and β-diversity in our mock community analysis compared to other data curation methods. This discrepancy resulted from the exclusion of ASVs perfectly matching (i.e., 100% identity and 100% query coverage) to insect species included in the mock community. This finding highlights the limitations of artefact identification methods that do not account for taxonomic assignments. However, our experimental design likely exacerbated LULU’s bias in two main ways. First, the use of multiple mock communities with slight variations of closely related species does not accurately reflect the natural diversity seen in environmental metabarcoding data, thereby complicating artefact identification based on co-occurrence patterns. Second, the default sequence similarity threshold of 84% employed in our study may be too low for the COI target gene (Hebert et al. 2003; Pentinsaari et al. 2016; Posada-López et al. 2023). Although LULU was developed for the ITS gene region, where an 84% threshold is generally appropriate (Sharma and Thakar 2024), the authors recommend adjusting the sequence similarity threshold for different target genes (Frøslev et al. 2017). Our meta-analysis indicates that few studies have adjusted this threshold and thus we followed the common practice in our experiment, though modifications to this parameter could potentially yield better results. Although our experimental design potentially exacerbated the bias associated with identifying artefacts based on co-occurrence patterns of similar sequences, our results were replicated within the reanalysis of air eDNA data from Lynggaard et al. (2022). The initial publication included a LULU data curation step, though reanalysis with *tombRaider* saw a large increase in the number of detected Copenhagen Zoo animals. These novel detections of zoo animals, however, underscore the enhanced potential of air eDNA biomonitoring, suggesting even greater utility than previously reported in this landmark study (Lynggaard et al. 2022).

Applying an abundance threshold to exclude OTUs/ASVs with low read counts resulted in the second greatest deviation from expected diversity metrics among all data curated methods evaluated. Unlike artefact identification based on co-occurrence patterns of similar sequences, the abundance threshold method removed fewer ASVs that were assigned to mock community species with a perfect match to the reference database (i.e., those with 100% identity and 100% query coverage). Nonetheless, this approach was also less effective at removing sequences flagged as artefacts in our study, despite our experimental design potentially favouring this data curation method. This favourable bias arises because the starting DNA concentration of the target gene across mock community species is likely less variable compared to the typical variation found in environmental samples (Hilário et al. 2023), which may result in fewer species with spurious read counts. The removal of ASVs assigned perfectly to mock community species suggests that setting an abundance threshold might be problematic, as read abundance can be influenced by amplification bias due to variability within the primer-binding regions among insect species (Elbrecht and Leese 2015, 2017; Loos and Nijland 2021). Conversely, the increased retention of artefacts likely reflects the creation of PCR errors during the early stages of amplification (Pienaar et al. 2006) and the potential presence of NUMTs (Andújar et al. 2021), which would be present in higher copy numbers than the threshold value (Pienaar et al. 2006; Andújar et al. 2021). Although increasing the threshold might eliminate more artefacts, such an approach also raises the risk of discarding genuine biological entities (Krehenwinkel et al. 2017; Liu et al. 2023).

Identification and removal of artefacts through the merging of ASVs with identical taxonomic assignments resulted in a β-diversity pattern that did not significantly differ from the known mock community composition and the taxon-dependent co-occurrence merging data curation method. However, a slight but statistically significant difference in phylogenetic α-diversity compared to the mock community was observed. These findings support the efficacy of artefact removal based solely on considering the taxonomic ID of ASVs. Nonetheless, caution is warranted when reference databases are incomplete for understudied geographical locations, taxonomic groups, or target gene regions (Weigand et al. 2019; Arranz et al. 2020; Leray et al. 2022), as this can lead to inaccurate taxonomic identification of OTUs/ASVs (Meiklejohn et al. 2019; Jeunen et al. 2022; Hilário et al. 2023; Keck et al. 2023). Furthermore, merging ASVs based on taxonomic ID is not suitable when haplotype data is crucial, limiting this approach for studies of population genetic structures using metabarcoding data (Andres et al. 2023; Couton et al. 2023; Liu et al. 2024).

We observed significant differences in diversity measurements based on the choice of data curation method employed, results in line with previous research (Frøslev et al. 2017; Andújar et al. 2021). Hence, comprehensive and accurate reporting is essential for the correct interpretation and replicability of metabarcoding studies. Our meta-analysis of 539 metabarcoding studies published in open-access journals during 2023, however, identified that current reporting for a majority of publications (437 publications; 81.08%) provide insufficient information to understand how sequencing data was processed. We advocate for improvements on data reporting standards as an essential component for advancing metabarcoding based monitoring techniques. *tombRaider*’s log file promises to solve this issue when added to the supplementary information of a publication by reporting on all essential information of the data curation step post bioinformatic processing, including the executed code, a list of all parameters and thresholds, and information on which OTUs/ASVs were deemed an artefact.

*tombRaider* is an easy-to-use command line software program and R package capable of accurately identifying and removing artefacts and pseudogenes from metabarcoding data. The incorporated modular algorithm provides users the flexibility the choice of which criteria are assessed in the identification of artefacts, while also facilitating the incorporation of novel criteria in the future upon advancements in the research field. The log output file, on the other hand, is generated to aid in standardised reporting standards for this emerging field, thereby enabling correct interpretation and replicability of metabarcoding results. Combined with sampling innovations (Mariani et al. 2019; Bessey et al. 2021; Hendricks et al. 2023; Jeunen et al. 2024), laboratory automation (Murray et al. 2015), and data curation post bioinformatic processing with *tombRaider*, eDNA metabarcoding now holds the potential for accurate, large-scale biomonitoring across the tree of life in a non-invasive, cost-effective, and time-efficient manner (Stat et al. 2017; Bowers et al. 2021; Takahashi et al. 2023).

## Supporting information

Supplement_4_tombRaider_log.txt

Supplementary_Table_1_tombRaider_meta_analysis.xlsx

## 5 ACKNOWLEDGEMENTS

A New Zealand Royal Society Te Apārangi Marsden Fast-Start (MFP-UOO2116) supported much of this project, as well as internal funding from the Marine Sciences and Anatomy departments within the University of Otago. We especially thank the authors of the three published datasets used to validate *tombRaider* for making their sequencing data freely available and easily accessible. Furthermore, we thank Dr Kristine Bohmann and Dr Christina Lynggaard for providing the full animal inventory list of the Copenhagen Zoo during eDNA sampling. We thank Prof. Stefano Mariani and Sadie Mills for their stimulating discussions on taxonomy and taxonomic assignment.

## 6 AUTHOR CONTRIBUTIONS

GJJ, KF, and NG designed the experiment. GJJ wrote the code for *tombRaider*. ED helped with debugging *tombRaider*’s code. QM and MT wrote the R package of *tombRaider*. KF led the meta-analysis, with help from GJJ, ED, GK, and AM. GJJ and KF wrote the manuscript, with significant input from ED, QM, ML, and NG. All authors contributed to the writing of the manuscript.

## 7 DATA AVAILABILITY STATEMENT

The *tombRaider* software program is open source and available on GitHub (https://github.com/gjeunen/tombRaider; https://github.com/gjeunen/tombRaider_R). Information on all papers included in the meta-analysis can be found in Supplement 1. Bioinformatic and R scripts used for data analysis can be found on GitHub (https://github.com/gjeunen/tombRaider_supplemental_files) and a detailed description is provided in Supplement 2, Supplement 3, and Supplement 4. A list of Copenhagen Zoo animals held during air eDNA sampling can be found in Supplement 5. Previously published data was used for the validation experiments, with links to raw sequencing files and metadata available in the respective publications (Braukmann et al. 2019; Weitemier et al. 2021; Lynggaard et al. 2022).

