## Supplemental Files for "*tombRaider* – improved species and haplotype recovery from metabarcoding data through artefact and pseudogene exclusion"

### Supplement 1: Meta-analysis

A systematic literature search was conducted to identify publications using metabarcoding methodologies. Peer-reviewed manuscripts published between the 1^st^ of January 2023 and the 10^th^ of November 2023 containing “metabarcoding” in the tile, abstract, or keywords were searched using the Scopus citation database on the 10^th^ of November 2023. We restricted our search to open-access journal articles to limit the number of publications and excluded conference proceedings, book chapters, and commentaries/notes. We, also, limited our search to articles written in English. We randomly sub-selected 600 publications to further reduce the number of included articles. We screened all publications individually to determine if methodologies employed metabarcoding techniques using the Illumina sequencing platform. Publications not meeting all requirements were excluded, with 539 publications remaining for the meta-analysis. This list of 539 publications can be found in Supplementary Table 1.1 (file name: “Supplementary_Table_1_tombRaider_meta_analysis.xlsx”). Contained within this file are also the metadata fields to create the alluvial diagram (Figure 1 in the main manuscript). The alluvial diagram was created using RAWGraphs *v* 2.0 (Mauri et al. 2017).

### Supplement 2: mock community data analysis

To investigate the impact of various data curation strategies post bioinformatic processing on diversity metrics, we conducted a comparative experiment based on a previously-published mock community data set (Braukmann et al. 2019). The terrestrial arthropod mock communities were comprised of 367 unique taxa, each constructed with different body segments (abdomen, bulk leg, composite leg) and slightly altered species compositions. Illumina sequencing data was downloaded from NCBI’s Short Read Archive (SRP158933) using the `fastq-dump` function within the SRA toolkit. As the forward and reverse reads were concatenated, VSEARCH’s *v* 2.16.0 (Rognes et al. 2016) `fastq_filter` function was utilised to split the file into an R1 and R2 fastq file. Reads were merged using the `fastq_mergepairs` function in VSEARCH using default settings. To remove primer sequences, cutadapt *v* 4.4 (Martin 2011) was employed, allowing for no insertions and deletions, as well as a maximum of two errors in the primer-binding regions. Remaining sequences were further filtered on quality using the `fastq_filter` function in VSEARCH, based on a maximum expected error of 1.0, a minimum length of 400 bp, a maximum length of 415 bp, and a threshold of zero ambiguous base calls within the amplicon. High quality amplicons were dereplicated using the `derep_fulllength` function in VSEARCH with default settings. Unique sequences were denoised using the UNOISE2 algorithm (Edgar 2016) implemented in VSEARCH (function: `cluster_unoise`) with default settings. Afterwards, chimeric sequences were removed using the UCHIME3 algorithm (Edgar et al. 2011) implemented in VSEARCH (function: `uchime3_denovo`) with default settings to generate a final list of ASVs. Finally, an ASV table was generated using the `usearch_global` function in VSEARCH. Since the ASV sequences consisted of a ~410 bp fragment of the cytochrome c oxidase subunit I gene, we assigned a taxonomic ID to each ASV through BOLDigger (Buchner and Leese 2020) with default settings. The resulting files were processed using a variety of data curation approaches to investigate the impact on the obtained diversity metrics, including: unfiltered (no data curation post denoising), taxon-dependent co-occurrence merging (criteria evaluated: taxonomic ID, sequence similarity, co-occurrence pattern), taxonomic ID merging (criteria evaluated: taxonomic ID), taxon-independent co-occurrence merging (criteria evaluated: sequence similarity, co-occurrence pattern), and abundance filtering whereby a detected was set to 0 if the read count was lower than 0.01% of the total number of reads within the sample. Except for abundance filtering, all data curation was conducted within *tombRaider v* 0.1.7. Differences with the known mock community were assessed through phylogenetic α- (Faith’s PD) and β- diversity measures (unweighted UniFrac distance). To generate the phylogenetic tree, all ASVs and amplicons of mock community species were aligned using the DECIPHER *v* 2.28.0 R package (Wright 2016) and exported in nexus format. The Bayesian phylogenetic tree was generated using BEAST *v* 2.7.6 (Bouckaert et al. 2019). Phylogenetic tree construction was performed with a Markov Chain Monte Carlo (MCMC) chain length of 10^8^ iterations, sampling trees every 1000 iterations. Convergence of the MCMC chains and effective sampling size was checked using TRACER *v* 1.7.2 (Rambaut et al. 2018). The maximum credibility tree from the posterior sample of phylogenetic time-trees with a burn-in percentage of 85% was identified through TreeAnnotator *v* 2.7.6 (Bouckaert et al. 2019) and used for subsequent analyses. The phylogenetic tree was imported into R using the ggtree *v* 3.10.0 R package, with identification for each data curation method visualized using the gheatmap function. Faith’s PD was calculated in R to investigate differences in phylogenetic α-diversity between data curation methods. Differences were assessed through a one-way ANOVA, followed by the *post hoc* Tukey HSD test in R. To assess differences in phylogenetic β-diversity, data was imported into the phyloseq *v* 1.46.0 R package (McMurdie and Holmes 2013). A distance matrix based on unweighted UniFrac was used to visualize differences in phylogenetic β-diversity through Principal Coordinates Analysis (PCoA) ordination. All bioinformatic and statistical code can be found in the GitHub repo (<https://github.com/gjeunen/tombRaider_supplemental_files>) under heading “2. Supplement 2: mock community data analysis”.

### Supplement 3: air eDNA data analysis

To determine *tombRaider*’s capability in recovering relics from published metabarcoding data sets, i.e., species and haplotypes missed during initial analysis, we reanalysed Illumina sequencing data from the Copenhagen Zoo air eDNA project (Lynggaard et al. 2022). Sequencing data for two amplicons (Taylor 1996; Riaz et al. 2011) were downloaded from the University of Copenhagen Electronic Research Archive (ERDA). Paired-end reads were merged using the `fastq_mergepairs` function in VSEARCH *v* 2.16.0 (Rognes et al. 2016)with the `fastq_allowmergestagger` parameter enabled. Merged reads were demultiplexed using cutadapt *v* 4.4 (Martin 2011) allowing for no insertions and deletions, as well as a maximum of two errors in the barcode + primer-binding regions. Afterwards, data were imported into R and analysed using the DADA2 pipeline (Callahan et al. 2016) with default settings to create a single ASV table and ASV sequence file per primer set. Custom reference databases were generated for each primer set using CRABS *v* 0.1.8 (Jeunen et al. 2022) using default settings. Reference databases were exported in BLAST format to enable taxonomic assignment of ASVs using local blastn *v* 2.10.1+ searches (Altschul et al. 1990). Data curation post bioinformatic processing was conducted using *tombRaider* based on the taxon-dependent co-occurrence merging approach (criteria evaluated: taxonomic ID, sequence similarity, co-occurrence pattern). To determine the presence of relics in the air eDNA data set, we retained the species list for which an ASV matched perfectly to the reference database (i.e., 100% query cover and 100% percent identity) and subtracted the species identified in the original publication. The remaining species were mapped against the Copenhagen Zoo species list that were held during air eDNA sampling (Supplement 5). All bioinformatic and statistical code can be found in the GitHub repo (<https://github.com/gjeunen/tombRaider_supplemental_files>) under heading “3. Supplement 3: air eDNA data analysis”.

### Supplement 4: salmon haplotype data analysis

To investigate the potential of *tombRaider* to recover intra-specific variation from metabarcoding data, we reanalysed the NADH dehydrogenase 2 (ND2) dataset from(Weitemier et al. 2021). Metabarcoding data from three salmonids were analysed simultaneously, including *Oncorhynchus clarkii*, *O. kisutch*, and *O. tshawytscha*. Briefly, separate ASV tables for each species were obtained from the online Supplementary Files associated with the original publication (Weitemier et al. 2021). The separate ASV tables were merged into a single table and the ASV sequences were exported from the table and formatted in a two-line fasta file. A highly curated salmonid reference database was generated for the ND2 target gene using CRABS *v* 0.1.8 (Jeunen et al. 2022) prior to conducting a local blastn *v* 2.10.1+ search (Altschul et al. 1990) to assign a taxonomic ID to each ASV. Once a separate ASV table, ASV fasta file, and taxonomy assignment file were generated, *tombRaider* was executed to identify and remove artefacts using the `--use-accession-id` parameter. The *tombRaider* log file can be found in the supplemental file named “Supplement_4_tombRaider_log.txt”. To visualize the efficiency of *tombRaider* to identify true haplotypes, a Bayesian phylogenetic tree was generated using BEAST *v* 2.7.6 (Bouckaert et al. 2019) after aligning all available reference barcodes and ASVs using the `AlignSeqs function of the DECIPHER *v* 2.28.0 R package (Wright 2020). The phylogenetic tree construction was performed with a Markov Chain Monte Carlo (MCMC) chain length of 10^8 iterations, sampling trees every 1000 iterations. Convergence of the MCMC chains and effective sample size was checked using TRACER *v* 1.7.2 (Rambaut et al. 2018). The maximum credibility tree from the posterior sample of phylogenetic time-trees with a burn-in percentage of 85% was identified through TreeAnnotator *v* 2.7.6 (Bouckaert et al. 2019) and used for subsequent analyses. The final tree was read into R and visualized using the ggtree *v* 3.10.0 R package. Haplotype networks, on the other hand, were generated for each salmonid species independently using the pegas *v* 1.3 (Paradis 2010) and adegenet *v* 2.1.10 (Jombart 2008) R packages. All bioinformatic and statistical code can be found in the GitHub repo (<https://github.com/gjeunen/tombRaider_supplemental_files>) under heading “4. Supplement 4: salmon haplotype data analysis”.

### Supplement 5: List of Copenhagen Zoo animals

The Copenhagen Zoo held a total of 184 chordates and 57 invertebrates during air eDNA sample collection (Lynggaard et al. 2022). The chordates included 17 Amphibia, 67 Aves, 4 Actinopterygii, 63 Mammalia, and 33 Lepidosauria (Supplemental Table 5.1). The air eDNA extracts were amplified using two vertebrate primer sets (Taylor 1996; Riaz et al. 2011). Hence, invertebrate taxa could not be detected through metabarcoding. The initial publication (Lynggaard et al. 2022) incorporated an OTU clustering bioinformatic processing approach, followed by data curation through LULU (Frøslev et al. 2017), i.e., taxon-independent co-occurrence merging. Lynggaard et a. (2022) reported the detection of 39 (21.2%) zoo animals (Supplemental Table 5.1). Reanalysis with *tombRaider*, however, detected an additional 50 (27.17%) species, bringing the total of detected animals kept in the Copenhagen Zoo to 89 (47.83%), including 3 (17.65%) amphibians, 36 (53.73%) birds, 0 (0%) fish, 48 (76.2%) mammals, 2 (6.06%) reptiles, and 0 (0%) invertebrates (Supplemental Table 5.1).

Supplemental Table 5.1: List of all 241 Copenhagen Zoo animals held during the air eDNA sample collection experiment (Lynggaard et al. 2022). Taxa are split up into major taxonomic groups. Scientific and common names are provided in the first two columns. Air eDNA detection by the initial manuscript (bioinformatic processing by OTU clustering and data curation through taxon-independent co-occurrence merging, i.e., LULU (Frøslev et al. 2017)) and by reanalysis with tombRaider is indicated by “X” for each of the 241 Copenhagen Zoo animals.

| Scientific name | Common name | LULU detection | *tombRaider* detection |
| --- | --- | --- | --- |
| Mammalia | | | |
| *Sarcophilus harrisii* | Tasmanian devil |  | X |
| *Vombatus ursinus* | Coarse-haired wombat |  | X |
| *Macropus giganteus* | Eastern grey kangaroo | X | X |
| *Macropus rufogriseus* | Red-necked wallaby |  | X |
| *Echinops telfairi* | Lesser Madagascar hedgehog tenrec |  |  |
| *Elephas maximus* | Asian elephant | X | X |
| *Tolypeutes matacus* | Southern three-banded armadillo | X | X |
| *Choloepus didactylus* | Linne’s two-toed sloth | X | X |
| *Myrmecophaga tridactyla* | Giant anteater |  |  |
| *Tupaia belangeri* | Northern tree shrew |  | X |
| *Lemur catta* | Ring-tailed lemur | X | X |
| *Varecia variegata* | Black-and-white ruffed lemur |  |  |
| *Galago moholi* | Moholi bushbaby |  |  |
| *Cebuella pygmaea* | Pygmy marmoset |  |  |
| *Leontopithecus rosalia* | Golden lion tamarin | X | X |
| *Saguinus imperator* | Emperor tamarin |  |  |
| *Saguinus oedipus* | Cotton-top tamarin |  |  |
| *Saimiri boliviensis* | Black-capped squirrel monkey |  | X |
| *Pithecia pithecia* | White-faced saki |  | X |
| *Papio hamadryas* | Hamadryas baboon |  | X |
| *Hylobates lar* | Lar gibbon |  |  |
| *Pan troglodytes* | Chimpanzee |  | X |
| *Cynomys ludovicianus* | Black-tailed prairie dog |  |  |
| *Mus musculus* | House mouse | X | X |
| *Rattus norvegicus* | Brown rat | X | X |
| *Cavia porcellus* | Domestic guinea pig | X | X |
| *Dolichotis patagonum* | Patagonian mara |  | X |
| *Hydrochoerus hydrochaeris* | Capybara |  | X |
| *Oryctolagus cuniculus* | Domestic rabbit | X | X |
| *Rousettus aegyptiacus* | Egyptian fruit bat |  | X |
| *Caracal caracal* | Caracal |  | X |
| *Panthera leo* | Lion |  | X |
| *Panthera pardus orientalis* | Amur leopard |  |  |
| *Panthera tigris altaica* | Amur tiger |  |  |
| *Mungos mungo* | Banded mongoose | X | X |
| *Suricata suricatta* | Slender-tailed meerkat |  | X |
| *Canis lupus* | Gray wolf |  |  |
| *Vulpes lagopus* | Artic fox |  |  |
| *Ailuropoda melanoleuca* | Giant panda |  |  |
| *Ursus arctos* | Brown bear |  | X |
| *Ursus maritimus* | Polar bear |  | X |
| *Zalophus californianus* | California sea lion |  | X |
| *Phoca vitulina* | Harbor seal |  | X |
| *Aonyx cinereus* | Asian small-clawed otter |  |  |
| *Ailurus fulgens* | Red panda |  | X |
| *Equus caballus* | Horse | X | X |
| *Equus quagga* | Plains zebra | X | X |
| *Tapirus indicus* | Malayan tapir |  | X |
| *Ceratotherium simum* | White rhinoceros | X | X |
| *Sus scrofa* | Domestic pig | X | X |
| *Hippopotamus amphibius* | Hippopotamus |  | X |
| *Camelus bactrianus* | Domestic Bactrian camel |  | X |
| *Lama glama* | Llama |  | X |
| *Tragulus javanicus* | Java mouse-deer | X | X |
| *Rangifer tarandus* | Reindeer |  | X |
| *Giraffa camelopardalis* | Giraffe | X | X |
| *Okapia johnstoni* | Okapi | X | X |
| *Aepyceros melampus* | Impala | X | X |
| *Damaliscus pygargus* | Bontebok | X | X |
| *Bos taurus* | Domestic cow | X | X |
| *Capra hircus* | Pygmy goat | X | X |
| *Ovibos moschatus* | Muskox |  |  |
| *Cephalophus natalensis* | Red forest duiker | X | X |
| *Hippotragus niger* | Sable antelope | X | X |
| Aves | | | |
| *Struthio camelus* | Common ostrich | X | X |
| *Rhea americana* | Greater rhea |  | X |
| *Numida meleagris* | Helmeted guineafowl | X | X |
| *Rollulus rouloul* | Crested wood partridge |  | X |
| *Gallus gallus* | Domestic fowl | X | X |
| *Afropavo congensis* | Congo peacock |  | X |
| *Callonetta leucophrys* | Ringed teal |  |  |
| *Nettapus auritus* | African pygmy goose |  |  |
| *Mareca sibilatrix* | Chiloe wigeon |  |  |
| *Phoenicopterus ruber* | American flamingo |  | X |
| *Goura sclaterii* | Sclater’s crowned-pigeon | X | X |
| *Otidiphaps aruensis* | White-naped pheasant-pigeon |  |  |
| *Nesoenas mayeri* | Pink pigeon |  |  |
| *Tauraco erythrolophus* | Red-crested turaco |  |  |
| *Musophaga violacea* | Violet turaco |  | X |
| *Cariama cristata* | Red-legged seriema |  |  |
| *Zapornia flavirostra* | Black crake | X |  |
| *Spheniscus humboldti* | Humboldt penguin |  | X |
| *Ciconia ciconia* | White stork | X | X |
| *Eudocimus ruber* | Scarlet ibis |  | X |
| *Theristicus melanopis* | Black-faced ibis |  |  |
| *Scopus umbrette* | Hamerkop |  |  |
| *Pelecanus crispus* | Dalmatian pelican |  |  |
| *Himantopus Himantopus* | Black-winged stilt |  |  |
| *Larosterna inca* | Inca stern |  |  |
| *Uria aalge* | Common murre |  |  |
| *Alca torda* | Razorbill |  |  |
| *Fratercula arctica* | Atlantic puffin |  |  |
| *Bubo scandiacus* | Snowy owl |  |  |
| *Colius striatus* | Speckled mousebird | X | X |
| *Bycanistes buccinator* | Trumpeter hornbill |  |  |
| *Bucorvus abyssinicus* | Abyssinian ground hornbill |  | X |
| *Upupa epops* | Eurasian hoopoe | X | X |
| *Merops apiaster* | European bee-eater |  | X |
| *Ramphastos toco* | Toco toucan |  | X |
| *Trachyphonus erythrocephalus* | Red-and-yellow barbet |  |  |
| *Charmosyna stellae* | Stella’s lorikeet |  |  |
| *Nestor notabilis* | Kea | X | X |
| *Melopsittacus undulatus* | Budgerigar |  | X |
| *Loriculus galgulus* | Blue-crowned parrot |  |  |
| *Agapornis nigrigenis* | Black-cheeked lovebird |  | X |
| *Psittacus Erithacus* | Grey parrot | X | X |
| *Ara ararauna* | Blue-and-yellow macaw |  | X |
| *Amazona aestiva* | Blue-fronted amazon |  | X |
| *Amazona amazonica* | Orange-winged amazon |  | X |
| *Amazona auropaliata* | Yellow-naped amazon |  | X |
| *Amazona barbadensis* | Yellow-shouldered amazon |  | X |
| *Amazona leucocephala* | Cuban amazon |  | X |
| *Amazona ochrocephala* | Yellow-crowned amazon |  | X |
| *Cotinga cayana* | Spangled cotinga |  |  |
| *Pycnonotus jocosus* | Red-whiskered bulbul |  | X |
| *Irena puella* | Fairy bluebird |  |  |
| *Garrulax courtoisi* | Blue-crowned laughingthrush |  | X |
| *Cinnyricinclus leucogaster* | Violet-blacked starling |  |  |
| *Lamprotornis chalybaeus* | Blue-eared glossy starling |  |  |
| *Lamprotornis superbus* | Superb starling |  |  |
| *Leucopsar rothschildi* | Bali myna |  | X |
| *Kittacincla malabarica* | White-rumped shama |  |  |
| *Ploceus capensis* | Cape weaver |  |  |
| *Ploceus nigricollis* | Black-necked weaver | X | X |
| *Lonchura oryzivora* | Javan sparrow | X | X |
| *Cacicus cela* | Yellow-rumped cacique |  |  |
| *Paroaria dominicana* | Red-cowled cardinal |  |  |
| *Riaris Canora* | Cuban grassquit |  |  |
| *Volatinia jacarina* | Blue-black grassquit |  | X |
| *Ramphocelus bresilius* | Brazilian tanager |  |  |
| *Leiothrix lutea* | Red-billed leiothrix |  | X |
| Lepidosauria | | | |
| *Emys orbicularis* | European pond turtle |  |  |
| *Astrochelys radiata* | Radiated tortoise |  |  |
| *Chelonoidis carbonarius* | Red-footed tortoise |  |  |
| *Stigmochelys pardalis* | Leopard tortoise |  |  |
| *Testudo horsfieldii* | Central Asian tortoise |  |  |
| *Cuora amboinensis* | Southeast Asian box turtle |  |  |
| *Cuora flavomarginata* | Yellow-margined box turtle |  |  |
| *Carettochelys insculpta* | Fly River turtle |  |  |
| *Agama mutabilis* | Changeable agama |  |  |
| *Pogona vitticeps* | Inland bearded dragon |  |  |
| *Brachylophus fasciatus* | Lau banded iguana |  |  |
| *Iguana iguana* | Green iguana |  |  |
| *Phelsuma grandis* | Giant Madagascar day gecko |  |  |
| *Rhacodactylus auriculatus* | New Caledonia bumpy gecko |  |  |
| *Stenodactylus petrii* | Petrie’s gecko |  |  |
| *Eublepharis macularius* | Leopard gecko |  |  |
| *Eumeces schneideri* | African gold skink |  |  |
| *Tiliqua gerrardii* | Pink-tongued skink |  |  |
| *Dracaena guianensis* | Caiman lizard |  |  |
| *Varanus indicus* | Mangrove monitor |  |  |
| *Varanus salvator* | Water monitor |  |  |
| *Varanus varius* | Lace monitor |  |  |
| *Morelia viridis* | Green tree python |  |  |
| *Python regius* | Royal python |  |  |
| *Acrantophis dumerii* | Dumeril’s ground boa | X | X |
| *Eunectes murinus* | Green anaconda |  |  |
| *Chilabothrus subflavus* | Jamaican boa |  |  |
| *Elaphe schrencki* | Amur ratsnake |  | X |
| *Lampropeltis getula* | Common kingsnake |  |  |
| *Lampropeltis Triangulum* | Milksnake |  |  |
| *Lamprophis fuliginosus* | Brown house snake |  |  |
| *Crocodylus mindorensis* | Philippine crocodile |  |  |
| *Crocodylus suchus* | West African crocodile |  |  |
| Amphibia | | | |
| *Tylototriton shanjing* | Emperor newt |  |  |
| *Bufo viridis* | European green toad |  |  |
| *Duttaphrynus melanostictus* | Asian common toad | X | X |
| *Epidalea calamita* | Natterjack toad |  |  |
| *Dendrobates auratus* | Green and black poison frog |  |  |
| *Dendrobates leucomelas* | Yellow-headed poison frog |  |  |
| *Dendrobates tinctorius* | Dyeing poison frog |  |  |
| *Phyllobates terribilis* | Golden poison dart frog |  |  |
| *Phyllobates vittatus* | Golfodulcean poison dart frog |  |  |
| *Bombina bombina* | European fire-bellied toad |  |  |
| *Bombina variegata* | European yellow-bellied toad |  |  |
| *Trachycephalus resinifictrix* | Mission golden-eyed tree frog |  |  |
| *Dyscophus guineti* | Sambava tomato frog |  |  |
| *Pelobates fuscus* | European spadefoot toad |  | X |
| *Pelophylax lessonae* | Pool frog |  | X |
| *Polypedates dennysi* | Denny’s tree frog |  |  |
| *Theloderma corticale* | Tonkin bug-eyed frog |  |  |
| Actinopterygii | | | |
| *Metynnis argenteus* | Silver dollar |  |  |
| *Hemiancistrus dolichopterus* | Suckermouth catfish |  |  |
| *Brachyrhaphis roseni* | Cardinal brachy |  |  |
| *Astronotus ocellatus* | Tiger oscar |  |  |
| Invertebrata | | | |
| *Achatina achatina* | Giant Ghana snail |  |  |
| *Pandinus imerpator* | Common emperor scorpion |  |  |
| *Brachypelma smithi* | Red-kneed tarantula |  |  |
| *Lasiodora parahybana* | Brazilian salmon tarantula |  |  |
| *Poecilotheria metallica* | Gooty sapphire ornamental tarantula |  |  |
| *Coenobita clypeatus* | Land hermit crab |  |  |
| *Therea olegrandjeani* | Cockroach |  |  |
| *Gromphadorhina portentosa* | Madagascar hissing cockroach |  |  |
| *Blaberus craniifer* | Death scull cockroach |  |  |
| *Sphodromantis lineola* | Ghanian mantis |  |  |
| *Phaeophilacris bredoides* | Cave cricket |  |  |
| *Carausius morosus* | Walkingstick |  |  |
| *Extatosoma tiaratum* | Giant prickly stick insect |  |  |
| *Platymeris biguttata* | Two-spotted assassin bug |  |  |
| *Dynastes Hercules* | Western Hercules beetle |  |  |
| *Eudicella colmanti* | African flower beetle |  |  |
| *Pachnoda marginata* | Sun beetle |  |  |
| *Gnorimus nobilis* | Noble Chafer |  |  |
| *Zophobas morio* | Giant mealworm |  |  |
| *Battus polydamas* | Gold rim swallowtail |  |  |
| *Heraclides anchisiades* | Ruby-spotted swallowtail |  |  |
| *Papilio cresphontes* | Western giant swallowtail |  |  |
| *Papilio polyxenes* | Black swallowtail |  |  |
| *Heraclides thoas* | Giant swallowtail |  |  |
| *Parides arcas* | Arcas cattleheart butterfly |  |  |
| *Phoebis philea* | Orange barred sulphur |  |  |
| *Greta oto* | Glaswing butterfly |  |  |
| *Danaus plexippus* | Monarch butterfly |  |  |
| *Idea leuconoe* | Chinese kite butterfly |  |  |
| *Caligo atreus* | Yellow-edged giant owl butterfly |  |  |
| *Caligo eurilochus* | Forest giant owl |  |  |
| *Caligo Memnon* | Giant owl butterfly |  |  |
| *Eryphanis Polyxena* | Bamboo butterfly |  |  |
| *Opsiphanes tamarindi* | Owlet butterfly |  |  |
| *Morpho peleides* | Morpho butterfly |  |  |
| *Agraulis vanilla* | Passion butterfly |  |  |
| *Dryadula phaetusa* | Banded orange heliconian |  |  |
| *Heliconius charithonia* | Zebrawing butterfly |  |  |
| *Heliconius cydno* | Passionflower butterfly |  |  |
| *Heliconius doris* | Doris longwing |  |  |
| *Heliconis erato* | Small postman butterfly |  |  |
| *Heliconius hecale* | Golden helicon butterfly |  |  |
| *Heliconius hewitsoni* | Hewitson’s longwing |  |  |
| *Heliconius ismenius* | Banded zebrawing butterfly |  |  |
| *Heliconius Melpomene* | Postman butterfly |  |  |
| *Heliconius sapho* | Sapho longwing |  |  |
| *Heloconius sara* | Small blue Grecian butterfly |  |  |
| *Archaeoprepona demophon* | One-spotted prepone |  |  |
| *Catonephele numilia* | Stoplight catone |  |  |
| *Colobura dirce* | Zebra mosaic butterfly |  |  |
| *Hypna Clytemnestra* | Jazzy leafwing |  |  |
| *Siproeta epaphus* | Rusty-tipped page |  |  |
| *Siproeta stelenes* | Malachite butterfly |  |  |
| *Atta cephalotes* | Leafcutter ant |  |  |
| *Anadenobolus monilicornis* | Millipede |  |  |
| *Archispirostreptus gigas* | Giant African millipede |  |  |
| *Telodeinopus aoutii* | Millipede |  |  |
